# Convergent Evolution in Tumor Genomes Targets Functional Domains

**DOI:** 10.64898/2026.05.31.729131

**Authors:** Hai Chen, Li Liu

**Affiliations:** College of Health Solutions, Arizona State University, Phoenix, AZ 85004, USA; Biodesign Institute, Arizona State University, Tempe, AZ 85281, USA

## Abstract

Tumor evolution is shaped by selective pressures that repeatedly favor similar functional outcomes across genetically distinct cancers. While convergent evolution in cancer has been studied at the gene level, this work investigates selection on smaller functional units, namely protein domains. Using >9,500 primary tumor exomes from The Cancer Genome Atlas, we quantified selection strengths acting on missense and truncating mutations aggregated by protein domain. This analysis identified 818 domains under significant positive selection across tumor types. Notably, approximately half of these domains belonged to genes that would be difficult to implicate using conventional gene-centric approaches due to low mutational recurrence or mutations outside functionally critical regions. We classified positively selected domains by evolutionary antiquity. The most ancient domains trace back to pre-eukaryotes and are involved in core cellular processes (e.g., DNA mismatch repair and metabolism) and tend to accumulate the highest numbers of mutations. The majority of positively selected domains originated in early eukaryotes and are enriched for regulatory control and cellular organization, whereas metazoan-specific domains are primarily associated with signaling and cell–cell communication. These results suggest that cancer preferentially exploits deeply conserved biology, with regulatory complexity driving tumor adaptation, while recent evolutionary innovations are relatively fragile and dispensable. Collectively, these findings establish a domain-centered framework for understanding disease mechanisms and developing therapeutic strategies. By focusing on shared functional domains, this framework enables the identification of functionally convergent therapeutic targets and provides a new perspective for interpreting drug resistance, tumor recurrence, and relapse.

## Introduction

Cancer is an evolutionary process driven by somatic mutations that provide selective advantages to tumor cells [1, 2]. Identifying these driver mutations is essential for understanding tumorigenesis and developing targeted therapies [3]. Advances in cancer genome sequencing have led to the development of numerous computational methods for detecting cancer drivers from patterns of somatic mutations [4]. Most of these methods operate at one of two scales. Residue-level approaches identify recurrent mutation hotspots but may miss functionally equivalent mutations dispersed across multiple sites [5, 6]. Gene-level approaches identify genes exhibiting an excess of protein-altering mutations relative to neutral expectations [7-9]. However, neutral mutations outside functionally important regions may dilute the signal of selection. As a result, many cancer-associated mutations remain difficult to detect despite their biological significance.

Protein domains represent an intermediate scale between genes and individual residues. As evolutionarily conserved structural and functional units, they mediate critical cellular processes, including DNA repair, signal transduction, transcriptional regulation, protein– protein interactions, and tumor–microenvironment communication [10, 11]. Focusing analyses on these functional units will mitigate the attenuation of selection signals caused by mutations occurring in unrelated regions in the same gene. Furthermore, as the same domain may be present in multiple proteins, mutations can be aggregated across genes to increase statistical power and identify convergent positive selection acting on common molecular functions. Domain-centered analyses therefore provide a powerful framework for uncovering cancer driver mutations that may escape detection by conventional gene- or residue-level methods.

Previous studies have demonstrated the utility of domain-level analyses for identifying significantly mutated regions and recurrent mutation hotspots [12-15]. However, existing approaches largely rely on mutation density or positional clustering and typically focus on missense mutations alone, which may lead to biased inference because they do not explicitly account for local mutation-rate variation or the selective effects of different mutation classes. For example, recurrent mutations may arise simply because a domain contains numerous methylated CpG sites or residues such as arginine, whose codons frequently overlap CpG dinucleotides and are therefore particularly susceptible to mutation [16, 17]. Such patterns can mimic positive selection despite being driven primarily by underlying mutational processes.

We previously developed Genes Under Selection in Tumors (GUST), an evolutionary framework that quantifies somatic selection at the gene level while accounting for sequence context and variation in neutral mutation rates [18]. In this study, we extend this framework from genes to protein domains and introduce Domains Under Selection in Tumors (DUST). Using DUST, we constructed codon-aware multiple sequence alignments (MSAs) for 14,684 conserved domains in the human genome and mapped somatic mutations from 9,523 tumor exomes spanning 33 cancer types onto these domains. By using synonymous mutations as a neutral baseline, we estimated selection coefficients for missense and truncating mutations and identified domains under positive selection. Our analyses revealed extensive domain-level convergent evolution both within individual cancer types and across multiple cancer types. By increasing statistical power through the aggregation of mutations across shared functional domains, DUST enabled the discovery of positively selected domains associated with rare driver mutations that would likely be missed by conventional gene-level analyses. Furthermore, by examining the evolutionary antiquity of positively selected domains, we found that cancer-associated positive selection preferentially targets ancient and deeply conserved functional units. Together, these findings provide new insights into the evolutionary mechanisms of tumorigenesis and highlight potential therapeutic targets.

## Methods and Materials

### Constructing mRNA-based MSAs of CDDs

The NCBI Conserved Domain Database (CDD) (https://ftp.ncbi.nlm.nih.gov/pub/mmdb/cdd/) provides the consensus sequences of 64,571 conserved protein domains. Because analyses of somatic selection require codon-aware modeling, we constructed mRNA-based MSAs for each conserved domain. Specifically, we downloaded the GenBank files of 133,711 human proteins from the NCBI RefSeq database (last accessed May 2024). Using the R/genbankr package, we parsed the GenBank files to extract CDD domains annotated within each protein and retrieve the corresponding peptide sequences. For each CDD domain, we collected all human peptide sequences annotated with that domain and aligned each peptide to the consensus CDD sequence. MSAs were then generated by progressively merging pairwise alignments using the consensus sequence as the anchor [cite custom r-script in github]. Finally, the aligned peptide sequences were back-translated to their corresponding coding sequences while preserving codon boundaries and reading-frame structure, which generated mRNA-based MSAs.

### Mapping somatic mutations to CDDs

Annotated somatic mutation data were downloaded from the GDC Data Portal (last accessed May 2024), generated from The Cancer Genome Atlas (TCGA) whole-exome sequencing (WES) data processed using the MuTect2 workflow [19]. Hypermutated and hypomutated samples were identified using the 1.5 interquartile range rule and excluded from further analysis. Based on functional annotations, synonymous, missense, nonsense, frameshift and in-frame insertion/deletion (indel) mutations were retained for downstream analyses. Using mRNA-based multiple sequence alignments (MSAs) as a common coordinate system, mutations from different proteins were mapped to shared conserved domain regions.

### Estimating selection coefficients

We adapted the gene-level selection algorithm implemented in our previously developed GUST program [18] for domain-level analysis. This algorithm considers nine mutational rate categories corresponding to transversions (1: A→C or T→G, , 2: A→T or T→A, 3: C→A or G→T, 4: C→G or G→C), transitions between purines (5: A→G or T→C ), transitions between pyrimidines at non-CpG sties (6: C→T or G→A) and at CpG sites (7: CG→TG or GC→AT), insertions (8: ins), and deletions (9: del). For each CDD domain, observed mutations across the nine categories were modeled using multinomial distributions, with expected counts of synonymous, missense, nonsense, frameshifting, and in-frame indel mutations estimated through saturated mutagenesis. Using synonymous mutations as the neutral baseline, the model estimates selection coefficients for missense mutations (ω) and protein-truncating mutations (φ, **Eq. 1**).

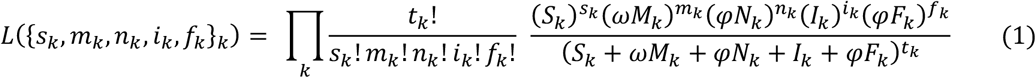

where *s*_*k*_, *m*_*k*_, *n*_*k*_ , *i*_*k*_ and *f*_*k*_ are the observed numbers of synonymous, missense, nonsense, in-frame indel and frameshifting indel mutations in the *k*^*th*^ rate category, respectively; *S*_*k*_, *M*_*k*_, *N*_*k*_ , *I*_*k*_ and *F*_*k*_ are the corresponding expected numbers of changes; and *t*_*k*_ = *s*_*k*_ + *m*_*k*_ + *n*_*k*_ + *i*_*k*_ + *f*_*k*_ is the total number of observed mutations. The parameters log(ω) and log(φ), constrained to the range [−5, 5], are estimated by maximum likelihood, where the sign and magnitude indicate the direction and strength of somatic selection, respectively, and values near 0 indicate neutral evolution.

### Evaluating statistical significance

To assess the statistical significance of positive selection for each CDD, we performed permutation-based simulations under a neutral model. For a CDD, given the observed counts of synonymous, missense, nonsense, frameshifting, and in-frame indel mutations, we randomly sampled the same numbers of mutations in each category from saturated mutation simulations and re-estimated the corresponding ω and φ values using the same likelihood framework. This procedure was repeated 1,000 times to generate null distributions of selection coefficients. The empirical *P* value was defined as the fraction of simulations producing ω or φ values greater than those estimated from the observed mutations.

### Annotating domains with evolutionary antiquity

We retrieved taxonomic annotations for each domain from the NCBI CDD database and the InterPro database [20]. Evolutionary age was inferred from the deepest taxonomic lineage in which a domain was detected, and domains were assigned to one of seven age categories: Root, Cellular Organisms, Eukaryota, Metazoa, Vertebrata, Mammalia, and Primates. In cases where CDD and InterPro provided conflicting age assignments, the CDD annotation was retained unless the InterPro annotation was supported by more than 1,000 sequences, in which case the InterPro assignment was used.

## Results

We mapped 14,684 CDD domains to human proteins and generated mRNA-based MSA for each domain. Somatic mutations identified in TCGA tumor samples were then mapped onto these domains, and selection coefficients were estimated.

### Distinguishing neutrally and positively selected domains

We analyzed 10,028 tumors across 33 cancer types. After excluding hyper- and hypo-mutated samples, the final dataset comprised 9,523 tumors, with sample sizes ranging from 40 cholangiocarcinoma (CHOL) tumors to 934 breast cancer (BRCA) tumors. Consistent with previous studies, tumor mutational burden (TMB) varied substantially across cancer types, ranging from a median of 6 mutations per exome in acute myeloid leukemia (LAML) to 381 mutations per exome in skin cutaneous melanoma (SKCM). Accordingly, the number of mutated CDD domains per tumor also varied widely, ranging from a median of 4 in LAML to 180 in SKCM (**Supplementary Table 1)**.

For each cancer type, we aggregated mutations occurring within the same domain across samples, resulting in 184,630 domain-cancer pairs, where the same domain appearing in different cancer types was counted separately. A vast majority (83.3%) of these pairs involved domains rarely mutated, with protein-altering mutations (missense and nonsense substitutions and indels) occurring in ≤5 samples. For the 30,757 domain-cancer pairs with >5 protein-altering mutations, we estimated selection coefficients.

Using synonymous mutations as the neutral baseline allowed us to separate mutation frequency from evidence of positive selection. We observed that frequently mutated domains are not necessarily under positive selection (**Fig. 1**). Notably, the C2H2 Zinc finger structural motif (CDD accession #: 275368) was the most frequently mutated domain in 31 out of the 33 cancer types analyzed, yet its selection coefficients (ω and φ) were consistently estimated to be near 0, suggesting neutral evolution.

**Figure 1.**
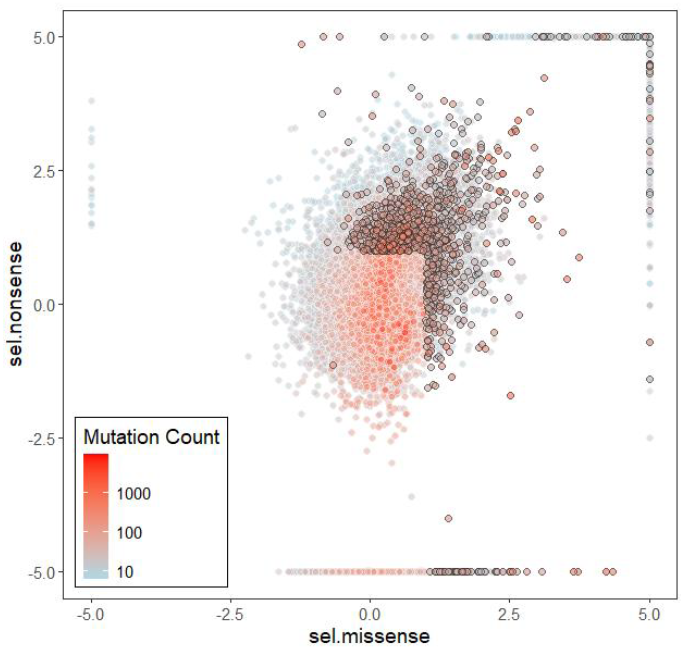
Scatterplot of the estimated selection coefficients, log(ω) and log(φ) for domains in the 30,757 domain-cancer pairs analyzed. Colors represent number of mutations observed.

Using an empirical *P* <0.1 threshold and requiring mutations in at >5% of tumors with estimated log(ω) or log(φ) values >1, we identified 1,190 domain-cancer pairs in which 818 unique protein domains were inferred to be under significant positive selection.

### Pancancer domain-level positive selection

The number of positively selected domains varied substantially across cancer types, ranging from none in adrenocortical carcinoma (ACC) and mesothelioma (MESO) to 408 in uterine corpus endometrial carcinoma (UCEC, **Fig. 2A**). Most positively selected domains were mutated in 5%–10% of samples, and their numbers declined sharply as the proportion of mutated samples increased. Only a small number of domains were mutated in more than 50% of samples. Some of these highly recurrent positively selected domains are found in well-known cancer drivers, including the p53 DNA-binding domain (CDD accession #: CDD:176262) in p53 tumor suppressor, the H_N_K_Ras_like domain (CDD:133338) in HRas, NRas, and KRas oncoproteins, the STKc_Raf domain (CDD:270964) in the Braf oncoprotein, the PTZ00435 domain (CDD:240417) in the Idh1 and Idh2 oncoproteins, and the Tryp_SPc domain (CDD:238113) in tyrosine kinases. Others are related to large protein families, such as the homeodomain (CDD:459649) found in homeobox transcription factors that regulate embryonic development, cell differentiation, and morphogenesis; the MutS domain (CDD:440019) present in DNA-binding enzymes that are involved in recognizing mispaired or unpaired DNA bases during mismatch repair and DNA recombination; and the WD40 repeat (CDD:441893), which act as rigid, versatile scaffolds to assemble multi-protein complexes and facilitate protein-protein or protein-DNA interactions.

**Figure 2.**
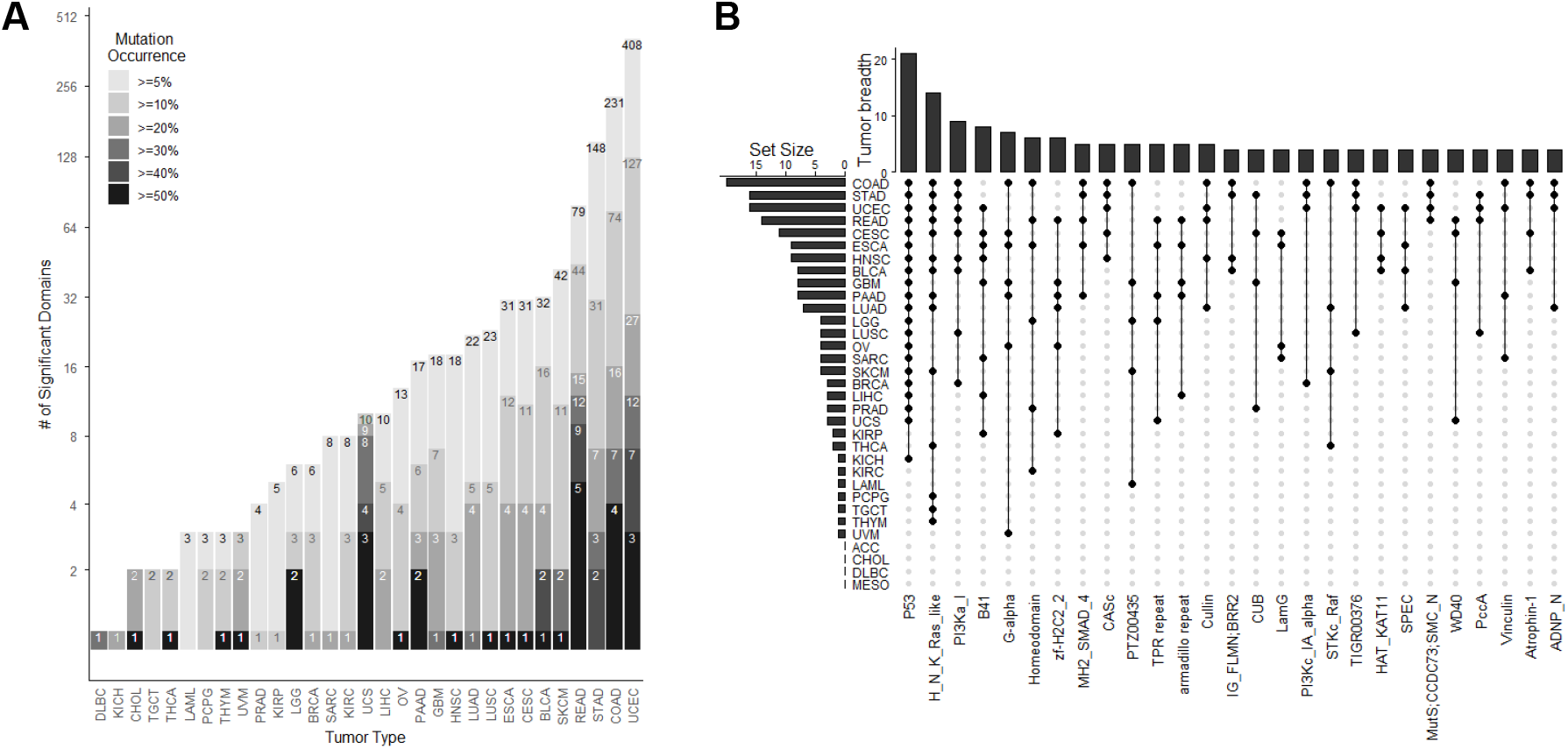
Positively selected domains across cancer types. (**A**) Stacked bar plot showing the number of positively selected domains in each cancer type, stratified by the percentage of samples harboring mutations within the domains. (**B**) Upset plot showing positively selected domains shared in at least 4 cancer types.

We observed a strong positive correlation between the number of positively selected domains in each cancer type and the total number of mutations sequenced (**Supplementary Fig. 1**). However, this correlation disappeared when restricting the analysis to domains mutated in at least 20% of samples. Therefore, we applied this threshold when identifying pan-cancer positively selected domains. Several positively selected domains were shared across multiple cancer types (**Fig. 2B**). The most prevalent was the p53 DNA-binding domain, which was positively selected in 21 cancer types. Other domains positively selected in more than five cancer types included the H_N_K_Ras_like, PI3Ka_I, B41, G-alpha, and Homeodomain domains. The H2C2 zinc finger domain was also positively selected in five cancer types. This domain is a functionally distinct variant of the canonical C2H2 zinc finger domain, which was under neutral selection in nearly all cancers. Specifically, H2C2 zinc finger domains are primarily associated with catalytic activity, multimerization, or cohesion machinery of genetic elements, whereas C2H2 zinc finger domains mainly function as DNA-binding motifs [21].

### Positively selected domains in specific cancer types

To identify cancer type–specific domains under positive selection, we required that a domain must be mutated in more than 20% of tumors and exhibit a strong selection coefficient (ω or φ >2) within a cancer type, but evolve under neutral selection in all other cancer types. Using these criteria, we identified three domains.

The first was the VHL beta domain (CDD:460360) in kidney renal clear cell carcinoma (KIRC), where the predominance of protein-truncating mutations suggests a tumor-suppressive role for this domain (**Fig. 3A**). The second was the GTF2I-like repeat domain in thymoma (THYM), in which a recurrent hotspot mutation pattern is consistent with oncogenic activity (**Fig. 3C**). The third was the U2 snRNP spliceosome subunit domain in uveal melanoma (UVM), which also exhibited a prominent hotspot mutation indicative of oncogenic function (**Fig. 3E**).

**Figure 3.**
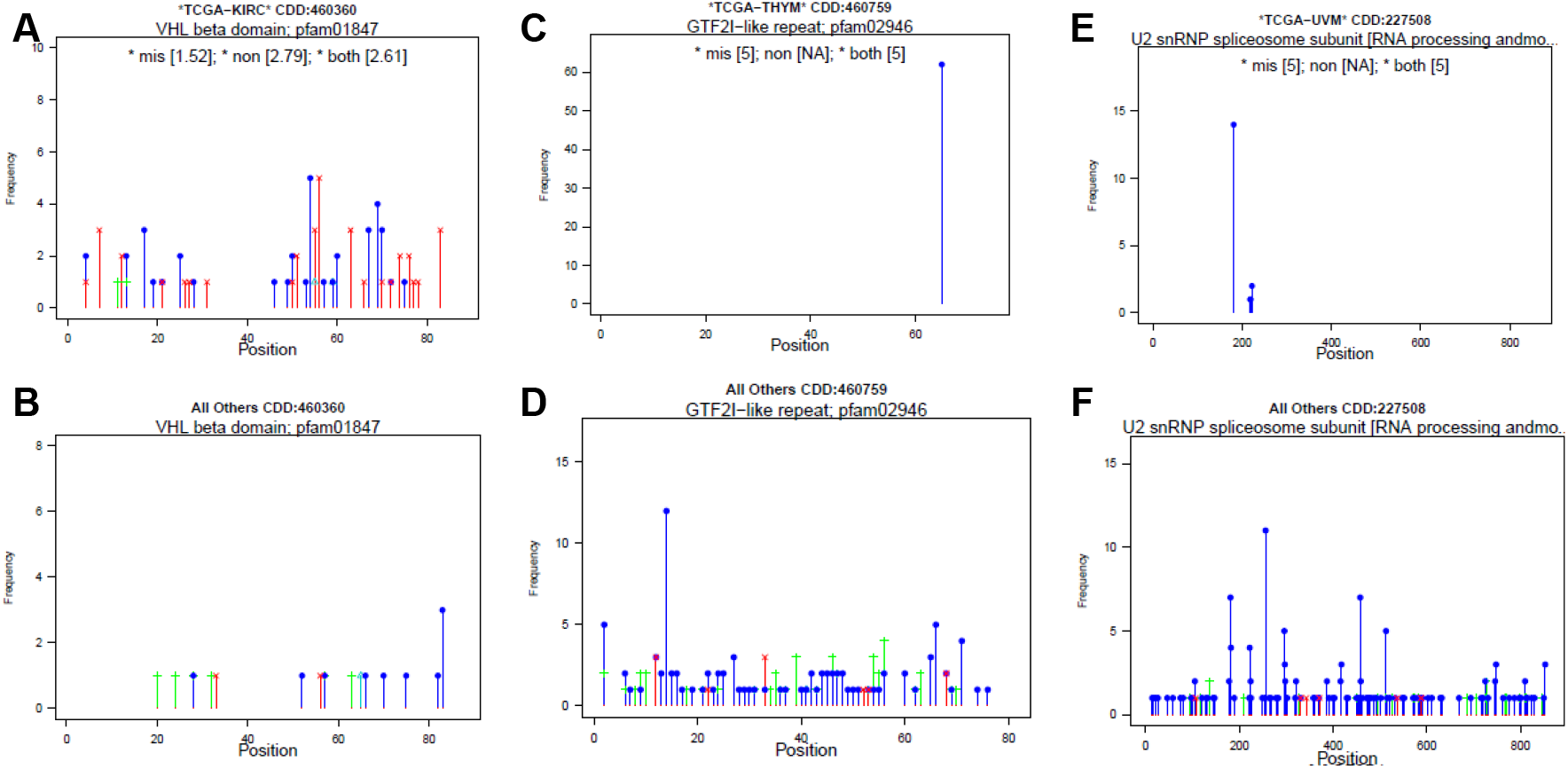
Domains under positive selection in a single cancer type. Positional distribution of synonymous (green), missense (blue), and truncating (red) mutations are shown. For cancer types in which a domain evolves neutrally, mutations were aggregated across all of them and displayed together for comparison. (**A-B**) VHL beta domain is positively selected only in KIRC. (**C-D**) GTF2I-like repeat is positively selected only in THYM. (**E-F**) U2 snRNP splicesome subunit is positively selected only in UVM.

These domains were rarely mutated in other cancer types. Even after aggregating mutations across all other cancer types, the distributions of protein-altering mutations closely resembled those of synonymous mutations (**Fig. 3B, D**, and **F**), consistent with neutral evolution. This pattern highlights the highly context-dependent nature of positive selection acting on specific protein domains in distinct cancer types.

### Mutations converged on positively selected domains

Among the CDD domains mapped to human proteins, 4,719 (32.1%) were present in multiple proteins. These shared domains provided an opportunity to examine whether positive selection acts convergently on the same functional domains across different genes.

We identified 558 domain-cancer pairs, involving 452 unique CDD domains, that exhibited evidence of convergent evolution across multiple genes. As expected, many involved members of the same protein family, such as the well-known Ras GTPase family (**Fig. 4A**) and isocitrate dehydrogenase family (**Fig. 4B**), in which the shared domain spans most of the protein sequence. We also observed convergent evolution in domains that constitute only a portion of the encoded proteins. For example, strong positive selection on the histone acetyl-transferase domain shared by chromatin regulators Crebbp & Ep300 (**Fig. 4C**) indicates that cancer-associated selection may converge on specific functional domains across distinct genes.

**Figure 4.**
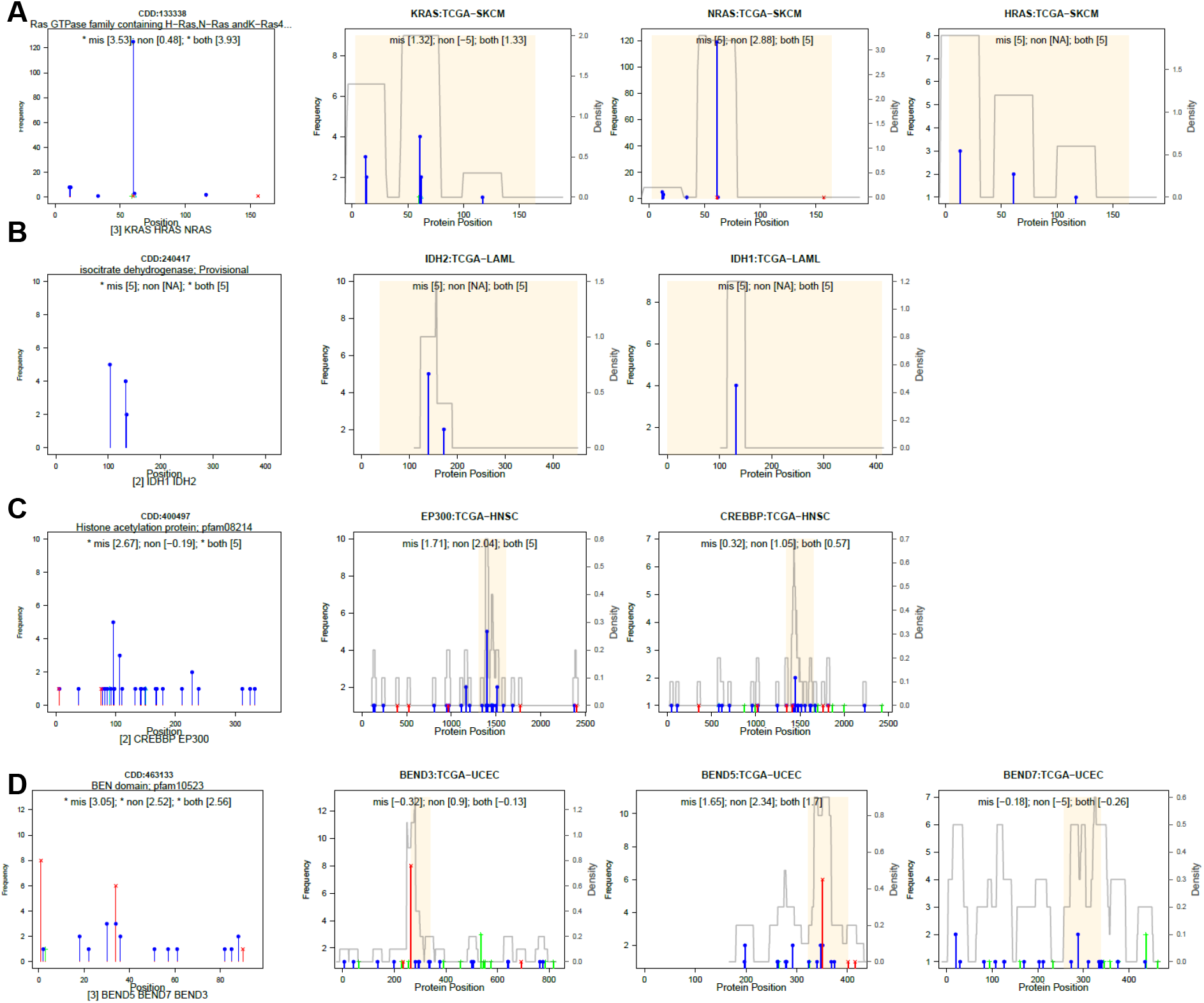
Convergent positive selection on shared domains across genes. Positional distributions of synonymous (green), missense (blue), and truncating (red) mutations are shown. In each panel, the leftmost plot shows mutations aggregated across all proteins containing the domain, and the remaining plots show individual proteins. Shaded regions indicate the locations of the shared domains within each protein. **(A)** The Ras GTPase domain shared by members of the Ras family. **(B)** The isocitrate dehydrogenase domain shared by IDH family proteins. **(C)** The histone acetyltransferase domain shared by Crebbp and Ep300. **(D)** The BEN domain shared by Bend3, Bend 5, and Bend7.

We also identified cases in which positive selection converged on a shared functional domain across multiple genes, while the regions outside the domain showed little evidence of selection. For example, the BEN domain shared by Bend3, Bend5, and Bend7 was recurrently targeted by missense and protein-truncating mutations, whereas synonymous mutations were concentrated in the remaining portions of the proteins (**Fig. 4D**). This pattern indicates that selection acts specifically on the BEN domain rather than on the proteins as a whole.

### Positive selection acting on rare mutations

Aggregating mutations across genes sharing the same functional domain enables the detection of positive selection that would otherwise be missed because of low mutation frequencies. This increased power arises in two distinct scenarios.

First, mutations may be individually rare across all genes containing a domain but become sufficiently abundant when pooled across genes to reveal a significant signal of positive selection. For example, in rectum adenocarcinoma (READ), *UEVLD, LDHB, LDHC, LDHAL6A*, and *LDHAL6B* genes each harbor only a few mutations. However, these genes share a subgroup of the L-lactate dehydrogenase domain (CDD:133429), and aggregating mutations across all five genes reveals strong evidence of positive selection (**Fig. 5A**). Similarly, eight genes containing the zinc-dependent metalloprotease domain (CDD:239797) collectively exhibit a strong signal of positive selection (**Fig. 5B**). Individually, these genes either harbor too few mutations to achieve statistical significance or contain synonymous mutations outside the domain that dilute the signal of selection when analyzed at the whole-gene level.

**Figure 5.**
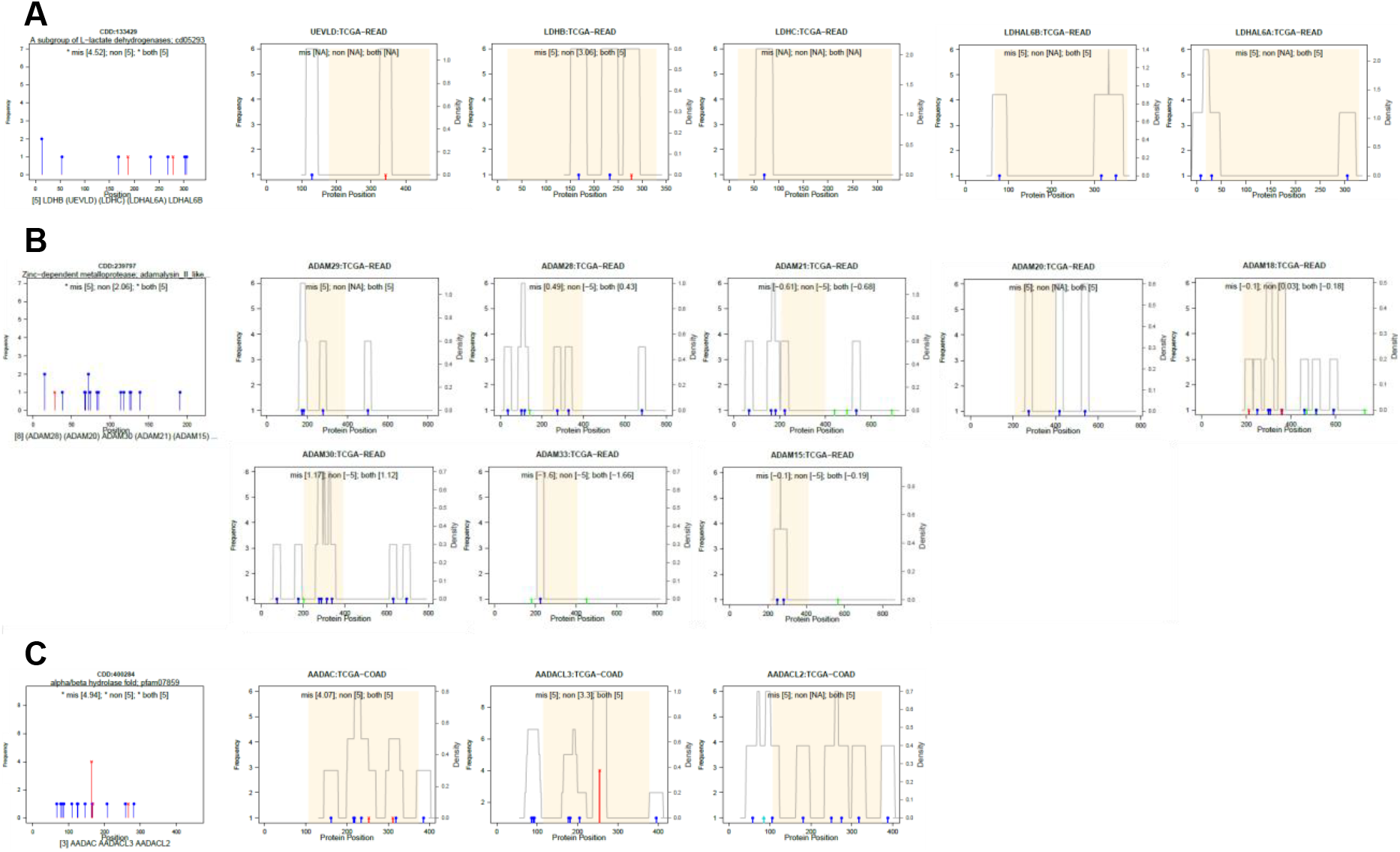
Domain-level aggregation reveals positive selection on rare mutations. Positional distributions of synonymous (green), missense (blue), and truncating (red) mutations are shown. In each panel, the leftmost plot shows mutations aggregated across all genes containing the domain, and the remaining plots show mutations in individual genes. Shaded regions indicate domain boundaries. (**A**) A subgroup of the L-lactate dehydrogenase domain in READ. (**B**) The zinc-dependent metalloprotease domain in READ. (**C**) The alpha/beta hydrolase fold domain in COAD.

Second, strong evidence of selection in one or a few genes can help uncover weaker but biologically consistent signals in other genes sharing the same domain. For example, in colon adenocarcinoma (COAD), *AADAC* and *AADACL3* are both under positive selection, and the alpha/beta hydrolase fold domain shared between them exhibits strong evidence of positive selection (**Fig. 5C**). A third gene, *AADACL2*, which also contains this domain, harbors only a small number of mutations and would not be identified as positively selected on its own.

Notably, although *AADACL2* contains both synonymous and missense mutations, the synonymous mutations occur outside the alpha/beta hydrolase fold domain, whereas all five mutations within the domain are missense substitutions. By leveraging the strong selection signal from *AADAC* and *AADACL3*, domain-level aggregation reveals a consistent pattern of positive selection in *AADACL2*, supporting its involvement in tumorigenesis despite limited mutational evidence at the gene level.

### Novel cancer-associated drivers

The domain-centered framework implemented in DUST enables the detection of cancer-associated drivers that may escape gene-level analyses because selection signals can be confined to specific functional regions. To evaluate the added value of domain-level analysis, we applied GUST to the same dataset, applied to same filter (selection coefficients > 1 and mutated in at least 5% of samples), and compared the resulting gene-level and domain-level selection signals.

We identified 517 domain–cancer pairs, involving 416 unique domains, that were inferred to be under positive selection by DUST even though the genes harboring these domains were classified as neutrally evolving by GUST. Applying a more stringent threshold (selection coefficient > 2 and mutated in at least 10% of tumors) to positively selected domains still yielded 13 domain–cancer pairs involving 13 unique domains.

A representative example is the dynein heavy chain N-terminal region 2 domain (CDD:462462) in uterine carcinosarcoma (UCS, **Fig. 6**). This domain is shared by multiple members of the dynein heavy chain family, including *DNAH5, DNAH8, DNAH9, DNAH17, DNAH2, DNAH7*, and *DYNC2H1*. When mutations were aggregated across all genes containing the domain, DUST inferred strong positive selection (log(ω)=3.49 and log(φ) =5.0). In contrast, none of the individual genes exhibited sufficient evidence of positive selection by GUST. Several genes contained only a few missense mutations within the domain, while others harbored synonymous mutations outside the domain that diluted the gene-level selection signal.

**Figure 6.**
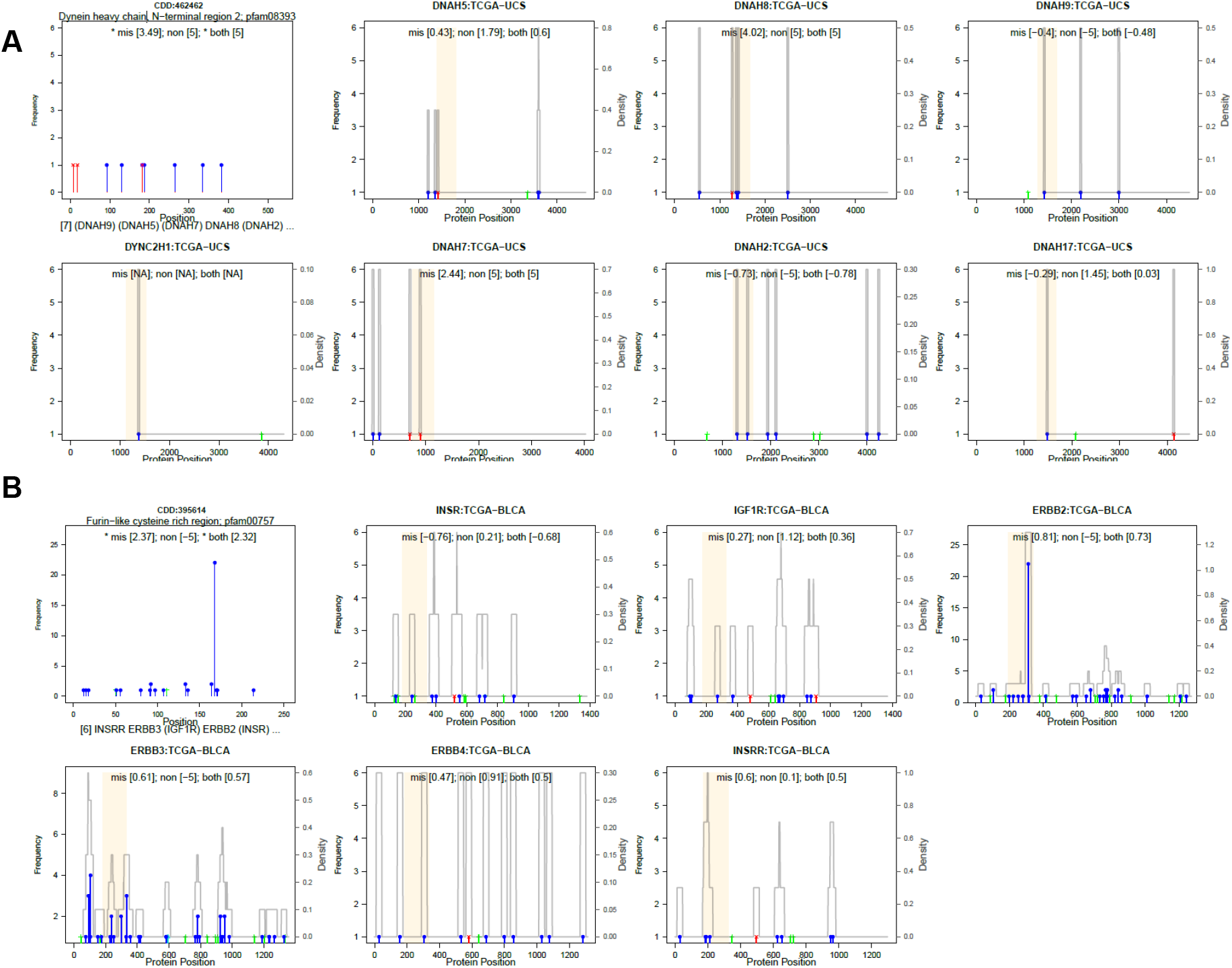
Novel cancer-associated domains. Positional distributions of synonymous (green), missense (blue), and truncating (red) mutations are shown. In each panel, the leftmost plot shows mutations aggregated across all genes containing the domain, and the remaining plots show mutations in individual genes. Shaded regions indicate domain boundaries. (**A**). Dynein heavy chain N-terminal region 2 domain. (**B**) Furin−like cysteine rich region.

Another example illustrates how domain-level analyses can separate functionally relevant mutational signals from neutral background variation to identify novel cancer-associated domains. The furin-like cysteine-rich domain (CDD:395614) is a 253-amino-acid domain shared by several receptor tyrosine kinases and growth factor receptors, including *IGF1R, ERBB2, ERBB3, ERBB4, INSR*, and *INSRR*, all of which encode proteins exceeding 1,200 amino acids in length. None of these genes exhibited significant evidence of positive selection when analyzed individually by GUST. However, inspection of the mutational distributions in bladder urothelial carcinoma(BLCA) revealed that protein-altering mutations were concentrated within the furin-like cysteine-rich domain, particularly in *ERBB2*, whereas synonymous mutations occurred predominantly outside the domain boundaries. By isolating this shared functional region from the remainder of the protein sequences, DUST was able to recover the underlying signal of positive selection and inferred strong positive selection for the domain (log(ω)=2.37, log(φ) =5.0).

### Convergent evolution targets domains from ancient to lineage-specific

Because cancer evolution acts on cellular functions that originated at different stages of biological evolution, we next investigated the evolutionary antiquity of positively selected domains. Specifically, if cancer preferentially targets fundamental cellular processes, domains under positive selection may be enriched among evolutionarily ancient protein domains. To test this hypothesis, we characterized the evolutionary antiquity of positively selected domains based on their taxonomic distributions.

We found that domains under positive selection were overwhelmingly evolutionarily ancient. The majority (68.5%, 815 of 1,190 domain–cancer pairs) originated in eukaryotes or earlier. For example, the G-alpha domain (CDD:206639), which exhibited strong positive selection (ω=4.9, φ=5.0) and was mutated in nearly all UVM tumors, can be traced back to the emergence of eukaryotes. In contrast, relatively few positively selected domains originated in vertebrates, mammals, or primates. Only two positively selected domains had a primate origin: Semenogelin in COAD and IgC1_MHC_Ia_HLA-B in COAD and STAD. All were mutated in fewer than 7% of tumors. This trend was even more pronounced among domains with higher mutation prevalence (>20% of tumors), where the proportion of ancient domains (originating in eukaryotes or earlier) was significantly higher than among less frequently mutated domains (84.5% vs. 66.8%, Fisher’s exact test, *P*=5×10^-5^, **Fig. 7A**).

**Figure 7.**
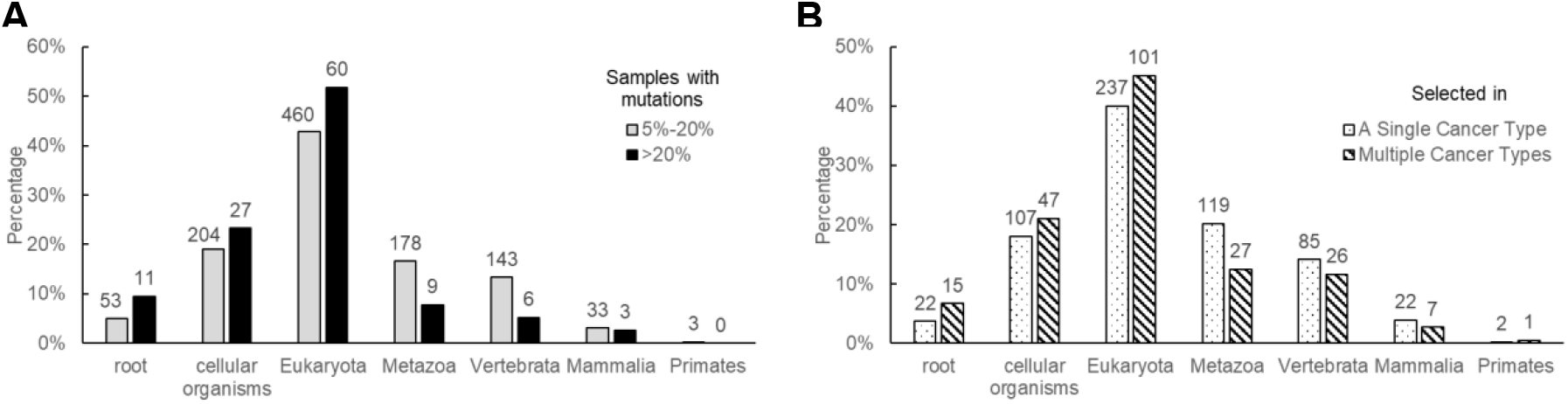
Evolutionary antiquity of positively selected domains. **(A)** Positively selected domains stratified by mutation prevalence. **(B)** Positively selected domains stratified by the breadth of cancer-type involvement. Within each stratum, bars represent the percentage of domains assigned to each evolutionary age category, and numbers above the bars indicate the corresponding domain counts.

Furthermore, domains under positive selection in multiple cancer types were significantly enriched for ancient domains compared with those under positive selection in only a single cancer type (72.8% vs. 61.7%, Fisher’s exact test *P*=0.003, **Fig. 7B**). The p53 DNA-binding domain, which was positively selected in 21 cancer types, and the H_N_K_Ras_like domain, which was positively selected in 14 cancer types, are both widely conserved across eukaryotes. The PI3Ka_I domain (CDD accession # 238444), which was positively selected in nine cancer types, is even more ancient, with homologs found throughout cellular organisms as well as viruses.

Together, these findings indicate that cancer-associated positive selection preferentially targets ancient and deeply conserved protein domains, particularly those that are recurrently mutated and shared across multiple cancer types.

## Discussions

Protein domains represent a fundamental organizational unit of biological function. By extending selection analysis from genes to conserved functional domains, we identified over 1,000 domain-cancer pairs under positive selection, demonstrating that protein domains constitute a powerful unit for studying somatic evolution in cancer.

A major advantage of domain-level analysis is its ability to detect convergent selection acting on shared functional regions across multiple genes. Classical examples include the Ras GTPase and IDH families, where mutations in homologous domains are known to produce similar functional consequences. However, we also observed convergence in domains that comprise only a small fraction of the encoded proteins, including the histone acetyltransferase domain shared by Crebbp and Ep300 and the BEN domain shared by members of the BEND family. Tumors harboring mutations in different genes but affecting the same functional domain may share common pathogenic mechanisms and potentially exhibit similar therapeutic responses and clinical outcomes. Domain-level analyses therefore provide a framework for grouping tumors according to disrupted molecular functions rather than mutated genes alone.

Domain-level aggregation also substantially increases statistical power to detect rare driver events, a longstanding challenge for both residue-level and gene-level analyses. Rare mutations are difficult to identify because individual sites or genes often harbor too few mutations to achieve statistical significance. In addition, gene-level analyses may dilute selection signals when positively selected functional regions are embedded within larger proteins that are otherwise evolving neutrally. By focusing on conserved functional domains, DUST separates mutations occurring in functionally important regions from those occurring in less constrained regions of the same protein. Furthermore, by aggregating mutations across proteins that share the same functional domain, DUST uncovered positive selection in domains that would have been missed by conventional gene-level approaches. This increased power arose in two scenarios: when mutations were individually rare across all genes containing a domain but collectively revealed a significant signal after aggregation, and when strong signals in one gene exposed weaker yet biologically consistent signals in related genes sharing the same domain. Domain-level analyses therefore complement existing driver discovery approaches by identifying functionally important selection signals hidden within the long tail of rare mutations.

Unlike genes, which can gain, lose, or rearrange domains during evolution, and individual sites, which are re too fine-grained to capture the evolutionary origins, protein domains often represent the fundamental structural and functional units upon which natural selection acts. Domain-level analyses thus provide a more direct connection between somatic evolution and evolutionary history than either gene-level or residue-level approaches. We found that domains under positive selection are overwhelmingly evolutionarily ancient. More than two-thirds originated in eukaryotes or earlier, and ancient domains were further enriched among highly recurrently mutated domains and domains shared across multiple cancer types. This pattern suggests that cancer evolution preferentially targets deeply conserved molecular functions that have remained essential throughout the history of multicellular life.

In summary, DUST establishes protein domains as an important intermediate scale for studying somatic evolution in cancer. By combining evolutionary modeling with domain-level aggregation, DUST distinguishes positive selection from mutation frequency, identifies convergent evolution across genes, improves the detection of rare driver events, and reveals novel candidate cancer-associated domains that are missed by gene-level analyses. These findings provide new insights into the functional architecture of cancer genomes and highlight the value of domain-centered approaches for cancer driver discovery and therapeutic development.

## Acknowledgements

The results <published or shown> here are based upon data generated by the TCGA Research Network: https://www.cancer.gov/tcga.

