## Supplementary Tables and Figures for "Convergent Evolution in Tumor Genomes Targets Functional Domains"

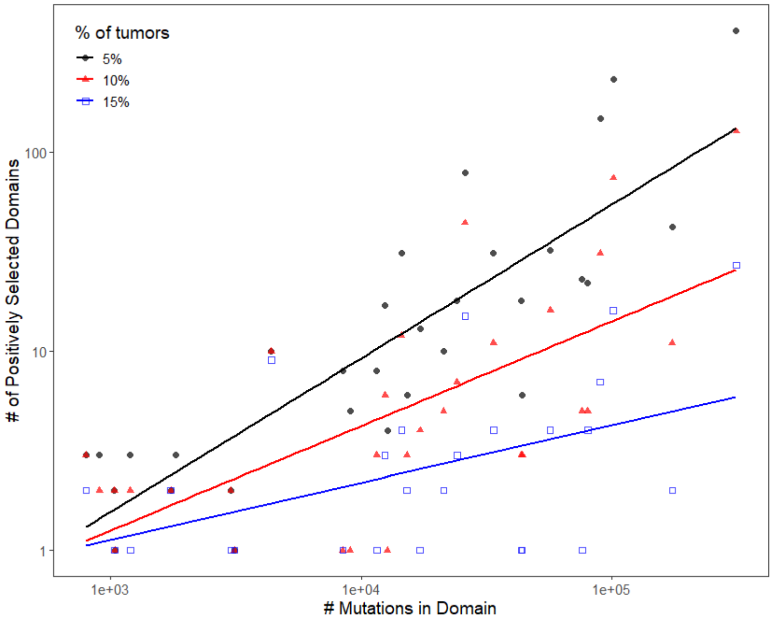


**Supplementary Figure 1.** Relationship between mutational burden and the number of positively selected domains across cancer types. Scatterplot shows the association between the total number of mutations mapped to protein domains in each cancer type and the number of domains inferred to be under positive selection. Positively selected domains were stratified by mutation-frequency thresholds of >5%, >10%, or >15% of tumors. Each point represents a cancer type, and solid lines indicate fitted linear regressions on the log-transformed scales.

**Supplementary Table 1**.

| Cancer Type | # of Tumor Samples | # of Mutations | # of Mutations in Domains | # of Mutated Domains | Median TMB | Median TMB in Domains | % of Mutations in Domains |
|---|---|---|---|---|---|---|---|
| **TCGA-ACC** | 85 | 7675 | 3979 | 2355 | 26 | 14 | 51.8% |
| **TCGA-BLCA** | 402 | 108137 | 56694 | 10363 | 184 | 92.5 | 52.4% |
| **TCGA-BRCA** | 934 | 82463 | 43832 | 9403 | 41 | 23 | 53.2% |
| **TCGA-CESC** | 274 | 64240 | 33649 | 8758 | 97 | 49.5 | 52.4% |
| **TCGA-CHOL** | 40 | 3344 | 1744 | 1280 | 37.5 | 21 | 52.2% |
| **TCGA-COAD** | 329 | 192488 | 101008 | 11725 | 119 | 63 | 52.5% |
| **TCGA-DLBC** | 46 | 6132 | 3132 | 1858 | 110 | 55 | 51.1% |
| **TCGA-ESCA** | 176 | 27636 | 14498 | 5529 | 114 | 61.5 | 52.5% |
| **TCGA-GBM** | 347 | 44055 | 24110 | 7172 | 54 | 30 | 54.7% |
| **TCGA-HNSC** | 486 | 81796 | 43451 | 9157 | 109 | 58 | 53.1% |
| **TCGA-KICH** | 63 | 2029 | 1045 | 831 | 17 | 9 | 51.5% |
| **TCGA-KIRC** | 357 | 21719 | 11538 | 4998 | 54 | 29 | 53.1% |
| **TCGA-KIRP** | 269 | 17541 | 9050 | 4513 | 62 | 33 | 51.6% |
| **TCGA-LAML** | 117 | 3381 | 1826 | 1324 | 6 | 4 | 54.0% |
| **TCGA-LGG** | 491 | 27762 | 15239 | 5646 | 28 | 16 | 54.9% |
| **TCGA-LIHC** | 347 | 41403 | 21333 | 6896 | 86 | 47 | 51.5% |
| **TCGA-LUAD** | 501 | 149706 | 80097 | 10248 | 199 | 103 | 53.5% |
| **TCGA-LUSC** | 462 | 142907 | 75968 | 10292 | 234 | 118 | 53.2% |
| **TCGA-MESO** | 75 | 2506 | 1288 | 994 | 29 | 15 | 51.4% |
| **TCGA-OV** | 383 | 32009 | 17192 | 6065 | 69 | 37 | 53.7% |
| **TCGA-PAAD** | 155 | 23195 | 12457 | 5250 | 37 | 21 | 53.7% |
| **TCGA-PCPG** | 176 | 1754 | 907 | 710 | 9 | 5 | 51.7% |
| **TCGA-PRAD** | 447 | 23061 | 12733 | 5253 | 26 | 15 | 55.2% |
| **TCGA-READ** | 127 | 48925 | 25966 | 7458 | 95.5 | 51.5 | 53.1% |
| **TCGA-SARC** | 220 | 16053 | 8448 | 3892 | 40 | 21 | 52.6% |
| **TCGA-SKCM** | 456 | 328760 | 173722 | 11545 | 381 | 180 | 52.8% |
| **TCGA-STAD** | 398 | 172142 | 89678 | 11261 | 123 | 67 | 52.1% |
| **TCGA-TGCT** | 122 | 2017 | 1039 | 823 | 14 | 8 | 51.5% |
| **TCGA-THCA** | 476 | 5365 | 3028 | 1795 | 9 | 5 | 56.4% |
| **TCGA-THYM** | 120 | 2241 | 1203 | 899 | 10 | 6 | 53.7% |
| **TCGA-UCEC** | 511 | 586201 | 312226 | 13175 | 81 | 44 | 53.3% |
| **TCGA-UCS** | 53 | 8202 | 4391 | 2602 | 52 | 31 | 53.5% |
| **TCGA-UVM** | 78 | 1405 | 802 | 560 | 12 | 7 | 57.1% |
